## Supplementary table 1 for "Heat Shock Factor Regulation of Antimicrobial Peptides Expression Suggests a Conserved Defense Mechanism Induced by Febrile Temperature in Arthropods"

**Supplementary table 1. Sequences of the primers used in this study.**

| **Primer** | **Sequence (5**'**-3**'**)** |
| --- | --- |

**Protein expression**

| nSWD-TRX-F | ATTAGAGCAAAGCCTGGCA |
| --- | --- |

| nSWD-TRX-R | TCAGAGGAGGTCAGGATTAGGCT |
| --- | --- |
| LvHSF1-PGEX-F | CGGGATCCTATGGCAGGAAGAAGCGGAGACAGCGACGAAGAATGCATGCCATTGAAGGATCT |
| LvHSF1-PGEX-R | ATAAGAATGCGGCCGCTCAATTCCCTCGTTTCTTCCC |

| **Quantitative PCR** | |
| --- | --- |
| EF-1α-F | TATGCTCCTTTTGGACGTTTTGC |
| EF-1α-R | CCTTTTCTGCGGCCTTGGTAG |
| β-actin-F | CACGAGACCACCTACAACTCCATC |
| β-actin-R | TCCTGCTTGCTGATCCACATCTG |
| LvHSF1-F | CCCAAGGACCTGCAATGGAT |
| LvHSF1-R | GCTCGACTCCTGACTCGTTT |
| nSWD-F | TGCGCTCGTGTTTTGTTCTG |
| nSWD-R | GGGATTTGAGATCGGGGCAT |
| RpL32-F | TGCTAAGCTGTCGCACAAATGG |
| RpL32-R | TGCGCTTGTTCGATCCGTAAC |
| DCV-F | GACACTGCCTTTGATTAG |
| DCV-R | CCCTCTGGGAACTAAATG |
| DmHSF1-F | TGGCCAGCTTCATAAGGCAA |
| DmHSF1-R | GCGATCAAAACGTAGTCCGC |
| DmAttacin-F | ACTGGCCAATGGTTTCGAGT |
| DmAttacin-R | GTTGCTGTGCGTCAAGGAAG |
| DmCecropin-F | AAGCTGGGTGGCTGAAGAAA |
| DmCecropin-R | CAGTCCCTGGATTGTGGCAT |
| **Absolute quantitative PCR** | |
| WSSV32678-F | TGTTTTCTGTATGTAATGCGTGTAGGT |
| WSSV32678-R | CCCACTCCATGGCCTTCA |
| TaqMan probe-WSSV32706 | CAAGTACCCAGGCCCAGTGTCATACGTT |
| **RNAi** | |
| dsGFP-F | ATGGTGAGCAAGGGCGAGGAG |
| dsGFP-R | TTACTTGTACAGCTCGTCCATGCC |
| T7-dsGFP-F | GGATCCTAATACGACTCACTATAGGATGGTGAGCAAGGGCGAGGAG |
| T7-dsGFP-R | GGATCCTAATACGACTCACTATAGGTTACTTGTACAGCTCGTCCATGCC |
| dsLvHSF1-1-F | TGAGGGATATGAAGCGCGAC |
| dsLvHSF1-1-R | GCTGTGGCTGTTGAAACTGG |
| dsLvHSF1-1-T7-F | GGATCCTAATACGACTCACTATTGAGGGATATGAAGCGCGAC |
| dsLvHSF1-1-T7-R | GGATCCTAATACGACTCACTATGCTGTGGCTGTTGAAACTGG |
| dsLvHSF1-2-F | ATCAACAAGTGGAGGGCAGG |
| dsLvHSF1-2-R | GGATGGGCTAATCTGTGGGG |
| dsLvHSF1-2-T7-F | GGATCCTAATACGACTCACTATATCAACAAGTGGAGGGCAGG |
| dsLvHSF1-2-T7-R | GGATCCTAATACGACTCACTATGGATGGGCTAATCTGTGGGG |
| dsnSWD-F | ATTAGAGCAAAGCCTGGCA |
| dsnSWD-R | TCAGAGGAGGTCAGGATTA |
| dsnSWD-T7-F | GGATCCTAATACGACTCACTATATTAGAGCAAAGCCTGGCA |
| dsnSWD-T7-R | GGATCCTAATACGACTCACTATTCAGAGGAGGTCAGGATTA |
| dsDmHSF1-F | ATGTGAAAGTCATGCGGGGT |
| dsDmHSF1-R | CGACGTTGTGCGAGCTAATG |
| dsDmHSF1-T7-F | GGATCCTAATACGACTCACTATAGGATGTGAAAGTCATGCGGGGT |
| dsDmHSF1-T7-R | GGATCCTAATACGACTCACTATAGGCGACGTTGTGCGAGCTAATG |
| dsDmAtta-F | ATGCAGAACACAAGCATCCTAATC |
| dsDmAtta-R | GCCGAAATGATGAGATAGACCC |
| dsDmAtta-T7-F | GGATCCTAATACGACTCACTATAGGATGCAGAACACAAGCATCCTAATC |
| dsDmAtta-T7-R | GGATCCTAATACGACTCACTATAGGGCCGAAATGATGAGATAGACCC |
| **Overexpression** | |
| pAc5.1a-LvHSF1-F | GGGGTACCATGCATGCCATTGAAGGATCT |
| pAc5.1a-LvHSF1-R | GGGGGCCCTCAATTCCCTCGTTTCTTCCC |
| pAc5.1a-DmHSF1-F | ATGTCCAGGTCGCGTTCATC |
| pAc5.1a-DmHSF1-R | TTACAACTCGTGACGTGGCG |
| **Dual-luciferase reporter assay** | |
| pGL3-nSWD-F | GGTTTTATCTATACCACTA |
| pGL3-nSWD-R | ATAAAAATACACATAATAT |
| pGL3-nSWD-M1-F | CCAAACGGCCAGGTTTGCTTCACTATAACA |
| pGL3-nSWD-M1-R | CCTGGCCGTTTGGAAAGTCCTAGTGGTA |
| pGL3-nSWD-M2-F | GGATTGGCTCGCCATTAGTGGCTGCGTG |
| pGL3-nSWD-M2-R | GGCGAGCCAATCCATCATGGAAATCCCAAC |
| pGL3-DmAtta-F | GCCATCAGGCCACCACCCATT |
| pGL3-DmAtta-R | GTTGCTGAACTGGATTGCTGGA |
| pGL3-DmCecA-F | GAAAAACAACTAAGTTACTA |
| pGL3-DmCecA-R | GGTGATATTTTCTTGATTTTTTC |
| pGL3-DmDef-F | TGACGCCAAAATGCAAGACA |
| pGL3-DmDef-R | CTTGGAATACAACTGGAGAGA |
| **EMSA** | |
| Bio-nSWD-Probe1-F | TACCACTAGGACTTTCTATAGAACCATCTTTGCTTCACTATAACA |
| Bio-nSWD-Probe1-R | TGTTATAGTGAAGCAAAGATGGTTCTATAGAAAGTCCTAGTGGTA |
| Unbio-nSWD-Probe1-F | TACCACTAGGACTTTCTATAGAACCATCTTTGCTTCACTATAACA |
| Unbio-nSWD-Probe1-R | TGTTATAGTGAAGCAAAGATGGTTCTATAGAAAGTCCTAGTGGTA |
| Mut-bio-nSWD-Probe1-F | TACCACTAGGACTTTCCAAACGGCCAGGTTTGCTTCACTATAACA |
| Mut-bio-nSWD-Probe1-R | TGTTATAGTGAAGCAAACCTGGCCGTTTGGAAAGTCCTAGTGGTA |
| Bio-nSWD-Probe2-F | GTTGGGATTTCCATGATGACTTTTCGAGGAATTAGTGGCTGCGTG |
| Bio-nSWD-Probe2-R | CACGCAGCCACTAATTCCTCGAAAAGTCATCATGGAAATCCCAAC |
| Unbio-nSWD-Probe2-F | GTTGGGATTTCCATGATGACTTTTCGAGGAATTAGTGGCTGCGTG |
| Unbio-nSWD-Probe2-R | CACGCAGCCACTAATTCCTCGAAAAGTCATCATGGAAATCCCAAC |
| Mut-bio-nSWD-Probe2-F | GTTGGGATTTCCATGATGGATTGGCTCGCCATTAGTGGCTGCGTG |
| Mut-bio-nSWD-Probe2-R | GTTGGGATTTCCATGATGGATTGGCTCGCCATTAGTGGCTGCGTG |
