## Supplementary table 2 for "Heat Shock Factor Regulation of Antimicrobial Peptides Expression Suggests a Conserved Defense Mechanism Induced by Febrile Temperature in Arthropods"

**Supplementary table 2. Heat shock proteins downregulated DEGs by transcriptome sequencing of dsGFP +WSSV (32 ℃) *vs*. dsLvHSF1 +WSSV (32 ℃).**

| **Gene lD** | **Description** | **log2(fc)** |
| --- | --- | --- |
| LOC113825261 | heat shock protein 21 [*Macrobrachium rosenbergii*] | -5.56 |
| LOC113806043 | heat shock protein 90 [*Litopenaeus vannamei*] | -2.53 |
| LOC113806042 | heat shock protein 90 [*Litopenaeus vannamei*] | -1.54 |
| LOC113825048 | heat shock protein 90 [*Scylla paramamosain*] | -1.33 |
| LOC113823664 | heat shock protein 90 [*Scylla paramamosain*] | -1.20 |
| LOC113802272 | heat shock protein [*Cherax destructor*] | -1.08 |
| LOC113815713 | Bip [*Litopenaeus vannamei*] | -1.16 |
| LOC113818927  LOC113830144 | Bip [*Litopenaeus vannamei*]  PREDICTED: dnaJ homolog subfamily B member 11-like [*Hyalella azteca*] | -1.13  -1.08 |
