## Supplementary table 3 for "Heat Shock Factor Regulation of Antimicrobial Peptides Expression Suggests a Conserved Defense Mechanism Induced by Febrile Temperature in Arthropods"

Partial promoter of AMPs was used in this study. The putative HSF1 binding sites in its promoter were shadowed

> Partial promoter region of SWD PP786678

TGTGTTTGTGTTGACTGAATAGTAAAGTGATATATATATATATATATATATATATATATATATATATATATATATATATATATATATATATATATATATATATATCTTAATGTTTACTCAAAAAAATCTTGTTTTTTTGGTCTCCAGAGCACTATAGAAAATATTTAACAAAATTATCATAATTTCCAAGAAAAATAAGCAAATAAATTTAGAGGTTTAGCATTTGCTAATTCTGTCCATTGCTATAGCAACAAAAATGTTTTTTTATTGGAAAAGTGTATGCAAGTAAACAAGCAGAAAACATAAATATTTATGAACAATATATGCATCATCATAACTTTGACGGGAGAAACGACAACATTTTGACAAGAAACGGTGAATTTGTTTTTCCTCGAGTACCCATGCTCCACTCTTTTTAACCGATACGAGTTAGTGATGACTGGTGACAGGCAAGTAAGATGATAATTATTTAACAGTTGGGTGGTTGGTGAGGATAATCATTTGAAATTCAACGGCTTTATTCACACACGCTCACCGGCCTCCCAATTGCTTCTTCGCCTCGACAAAAAAAAAAAAAAAAAAAAAAAAAAAAAAAATGGTTTTATCTATACCACTAGGACTTT**CTATAGAACCATC**TTTGCTTCACTATAACATTTTAAACGTGTGAATAGCCACTTATCTATGTTCCTAATAAAATATTCTGTGAAGTATCAGCCATTCATTTCAATTTATCCTCTGCCTATCTATCAGTCATTAAGATCTGTCCATAAATCTCTGTCCAACCACAGACAAGTAAACTCCATTTTTTCCTAACTTACGGTTTCGCAACAACGACCCTTTGCGCCATAACACCGGCTCCTCGGTGGATAAAAGGGCAGCGCTTGGCCCCTTGATGCTTCAGTTCAGATCCGTCCTACGTCGCTTGCTCATCCGGCACAGTTGGTTGGGATTTCCATGAT**GACTTTTCGAGGA**ATTAGTGGCTGCGTGGTGTGTGTCGCCGTGATACTTATGATGGTTGGTGTTTTGTATTTTTCGCGTTTTAGCATTAGTTTTTCATGTATTTATTCATGTGTAATGTATATTATGTGTATTTTTATGTTGTACATGTACGCGCGTGTCTGTCTGTCTGTACGAGTGTGCACGTAGATTCACATGTGCGTATGTCTACTTCTTAATCGATCAGGAATGTCATGAATCTATCAACCATGGATATAGGATCCACTGCACCTATTCTGATCCCCTTTCCTAACTCTGCCGGCGTTCGTGCTCTCCCCAGACGAGCACCGGACAGGCGGGTCCTACAGGCAGCCAGAAGCCAGGATCTTGCCCGTTATTCCCTGCTCCTCCTCCGCCTCACCTAGACGGGTAGGATATTTCTTCTTTGTTCTATTCTCTTTCTTATTCGTATTTTGTTATCATTATTCCTTTTTTTTTAATTCTTATGCCTATCCGTGTTCGTTTGGCTGAAGGCAGTAAAAGAGTAATAGGAAAAATATTTTTATGTCATTTTTCTACTGTTGGCACTATTTTACCATCAACGAAACTATTATTGTCGCAATTTTAATAGACAACTGTGGCCATAATTGTCTGTATCATTCATTAATGTCTTCGTTATCATGCCTTTAGACCACTGTTATCTTCACCCTCACCATCACTATCATTATCACTGTTATCGTTGCTGGCAATTTTTGTTATATATTTTTTGCTTCTATCTTTTTTATCATCACATCATGCAGCCGCAGATCTTGGAAAATCTTTATAAGCCGCTGTGAAGGGGGGGGGGGTGGTGACCCCAGTGAGACCGTGATGGTAATGATAGTGATAATGATGTATAATGCTAATTTTAATGATGATGGTGGTGGTATGACGGCGATAATGATGGTGATAATGATAATTTTGATAAAAACTATATTAATATCATTCCTACTACTACCACTAGTGCATTATTATTACTACTATCATTTCATAATCATCTTTATTATCATGATCGTTAT

> Partial promoter region of Attacin A (AttA) AY056895.1

GCCATCAGGCCACCACCCATTCTGCCCGCCTAAAGATGTGTGCATACCGCGGAGAAGTCATCCGATCAAATTTGTTTTGAAAAATCTTTATAAAAATTGTGAATTTTTTACT**TTCTGCAA**ACAGTAAGCAATAAACACACGAAAGACAGCAATTAATAATCTTCAATCAATTGTGACACAATGAGGGGTTCCCATCGCTTATCAGCGGTTTTTGTACCGAATCTGCTGAGCTCTAGAGCTGATAAGAAATATACTTGCTCAAAACAAAACCACAAAAGTCACGTTTGAGAGAAAAAAAGCCTAAACGAATTTAATTCGCGACTCATATGAATCACAAACC**TGTTCTAT**AGCACGTTCTTCTTAAATTCTAGCGAAATAACCAGATGGCTGCAAATCATAATGAATGGGTTTGTCCCCTAAAAAAAACGAACTGACAAGCCCCCTTATAAAACTTATTTATTAATAGATTAGTTCGTATATAATTGCATATGTAAATACTTTAA**ATAGAACA**AATTATTACGACATTTAAAAATATATATCCTGGTTTTTAAAAACAGGGTTTGAAAAAAATGTTTATAAGCTAATTACCTGGTAGATTATATTTTTCAGTGCAAACTTTTCGTTGACACTGCGGGTTAAAGTTTCCGCTCTCCTCTCTTTCGGCTCGCATCCTTTTTGCCCGCTCCTGCGCAGAAAATTCAATTAGGATTCGGCTGAAACTTCACTCAAATCCTGCCGCTCTCACCGTCGCTCTTTATTTCGCTCGCCTTCCCTTTCCGGCTCCCCCAGAATAATCCCTCCACGCAAAATAACGTATTGATAAAGCCTGACATCAAGTGAGAAATACGATAGAGAATCCCTTATAACTGTCAAATGCCGACCTGCGCAAGATGAGGATGCACTCCTCCATCAAGACACAAAAGAAACCACTTGGAGACGCTGACAGAGGTTCTGCGGCGAGGGTGAAACTGGACAATGCAGCAACAAGTGGCGTCAATGGGTCGCAAAAAAGGGGAGGTGATGAGGTCAAGTCGCACAGCCACACAAGCAAACAGCAGAAGTAAACCACCATCACATCTGAGCGGGGAATTTCGCTTTGATAAGGCATCCAGGCCGAGATCGGCAATCAGATGAATCATGTCAATCATCAGAAAAGTTCTTCCCCGCATCTTGAGGTATAAAACCGATGCATTGGACACCTTGAAACATCAGTCAGCTCCAGCAATCCAGTTCAGCAAC

> Partial promoter region of Cecropins A (CecA) AAF57025.1

GAAAAACAACTAAGTTACTAACGCAAGACTTTTAGTTAAGTTAGTTAATATAGACCGAGATGTATGTACATACATACCGCTTTCGCTTACAATAAAATGTTAAATAAGTTTTCAGATTCGTACGTGCTCAGTAAACAATTATTTTTTATTGTCATTTAATGCCTATTGAATTTTTCAAACTTAATTTAGTGCCTTTAGTAAAATATTGTAGTGATTCCCCTCGAAAAATACCACAAATTGGATGCGTTTATGTAAATAAATTGCCCTTGAGTGATAGAGTAAATTTGAATTTGACTGTCTTAGAAAG**ATAGAAA**GAGATCAATTCAAAATGCCAAAAGGATAGAGTTATTAAAGCTCTAATTCAAATTGGCCCAGAACCGTTTAAAGGATATTACAATTTGTAATTTACATATTTGGATTATAGCATTGAAATCCCCGAT**TGTTCCCT**AGATGTGCAGATGTGTGC**TTGGAATC**AGATCGGTTACCTTCAGTGTACTTTTCTCTGCAAAAATCCCCGTGCATGCCTTATCTGTCATTTTGTTTTTCAAGCTGGCTGTTCGCCTATAAAAGCTCTCGCCTTTTGTATCGCAGTCATCAGTCGCTCAGACCTCACTGCAATATCAATATCTTTAGCTTCTCCTAAGAAAAAATCAAGAAAATATCACC

> Partial promoter region of Defensin (Def) AAF58855.1

TGACGCCAAAATGCAAGACAAGACAACCTGGTCGATAACAAAGGTAAACAGGCAACGGCCAGCCAAGGAGCAGGGCAACTGAAAAAGCCTCGGTCGTGGGATTCGATCGGGATTGTCGGCTCAGCGCGACTGGGCTGAGAACCAGATGCAGATGCCGATACAGATATAGATACACGTACGCGCAGATACGGATTCAGATACAAGTACACCCGCCCCTGCCGCTGTATGCCCAACTAATCATTGTGTGATTCTTGTTTGTTTATTTGCCCGGCATTATGAAGAGACTTTTCGGTAGAAATTATTTATTGTCGCATGTGTTTATGTATCCGTAACCGAGTATCTCAGTTGCTTGAGCCAACTGTGTAGCTGTGTAGCTGTGAGTATAGCCCTTAAAGTGGCACCCAATCGGTCAGTTAGCTAGAAATTCAGATGATTAAATATGGATTCCCCTACATCAGCTAATTTCAACAGTTTGGGAGTAATAAAATCGAAATTGGATGCTACTAAAGGGCACATATTTACTTAGGCTTTTATCAACGTTGCATATATACAAATATCCTGCATATTTCGCAAACCAAAGATTCTTTCTCAAGTAAGGCCTAAACAATTTGAAATGGTTAATTTCGTAGATGTTGCTTTTTACAATTAACTTGTCAT**GTGGAATA**TACTTTACTGCCTAAAATTTAAGGCAGTTAAAATCCCTAGAAATGCAAATAACTTA**TTGCAGAA**ACGGGCTCTGTCGGCTGTATTTTGCTCTTATCTATGAAATATTGTCAATATTTTCCAGGCAAAGCACATGAAATAATGATCTAGACAACGGTTTCTCCCATTTGCAGTGAACTTAAAAATTAAAAACCCCCGAGACGTGTCTTCCTGCACAGAAAAAGAGACAATGGGAAGGTAAGTCACCGGGTGGGAGTCCCTGGGCCGAATCGATCAGCCCGTCGCATTGCTATATAAGCTCGGCGAAACCACAATCTGCAACAACAGTATCTCTCCAGTTG**TATTCCAA**G

> Partial promoter region of Metchnikowin (Mtk) AAF58139.1

ATTTAAAGGGTAGCGCCACGTTCAACCTCTTTTGCAGCCCCATTCTGCTGCGAGAAAACTAACAAAGTGCTCTAATCGAGCCAAGGGGCAATTTCTTGTGTTGCTGCAGCTGCACTTTGCACCTCCGCATCCGTGCACCCAAAAACCCGCTTTCTAGATGTTCTCATCATGCACTGAAAAAGAATCCAAATTTTTACAAGAAATAGTTTAAAATTAGGTAATGTGAAAGATATCGGCACACGGACAGGCCGAATTATCTGTTGTAAAACTAGCTGCTCAGTTATTAAAAACATTTGTAGTTGCTGACGTTTCCATACAGAGACTAATTTTATTTTCACGGACAGGGGTTTTCCGCTTTAATTGCTTCATTTTTGTTGCTTTATTGCGTGTATATTGCCCCACAAAACAGATATAAATCATTCGCGCATATCGTAAATGTTGGTAGAAAATGACAAACAAGAGAAAAAAGATGATTAAAAGCTTCAAGACAATCCTCTATAGGATCTGATTAAATATGAATATTTTATTTATTAGTTTTCTTTCTGTGTACGGCTTAGAAGGCAGAAGCTGCGAGGGGCGTAGGGCAGTGGGCGTGGCTCCGTGTTGACGCATGTTGACTATGCCTTTGAATGGCTGCCGTGGTTGTCGGTGGGTAATTTGCAATGCAGAAAAACCAACAGGGCGCTAAAAAGGAGAGTGTTTTCGTGGGAGGTGGAGATGGTCACTGGGGGCAACATAAATATTCAGCGAGAAACGTCATATTTACATTTAGTCTAGGCTGATAATCCGGGACCGTGGGAAGTCCCCTTTGGGTGGTGCTGGCTGGGTTCCCCTGGCCACAATCGGTTATCTGCCCCCGGCTGACACTTGCCCGTCATTCATTCGGCTGCTTATCGCAGAAGCTCAAATAAAAAGTCCCCAATCTGCGACTCGTTTGTCTGGGACTGAGCTATAAAAGCCTCACCATCTCAACGCTCAAAGCATCAATCAATTCCCGCCACCGAGCTAAG

> Partial promoter region of Drosomycin (Drs) AJ885064.1

CAATGAAAGTGATAATACGAATTGACCATGTAGCAATTTGTTTTGTGTCTATAGACTGAATTTTTTCCCCTACATTAATCAAAATTATATTTTTATATTTTATTTTGAGTTTACTTGGGTTTTTCATGAAATTAAAGATTAACCTGGGGTTTTTACAATCCATTACGATAGGCTTTCTGTTCATTGCATCTATAGCCTCTGTACTTTTCCGTGCATTCTTAAAGCAAAAGCATACATGTATATCTTCAATTCAAGTATATCTGCAATTTAGTTTGCTTATCTGGAGTCGCTGTATCCCGACCTATCCAAACTCGCGTCCCAGTCAAAGGTAAACCATTTTTATTTAGTTCCCAGCCTCTGGTATTTGTTGTTATTTATCGGTGACTTTTTGAAATTTATTCAATTAATTTCAGTCTTTGTGCTCTTGATAACACGATTTCCTCGTTATTCATTTAGTTTGGGTTTAACCAAAACCCTTTAGCGAATCATTTTTGCTGGACAGTCCAGTTGAATTCGGTATTCTACACACAAAGCTCATCTTACAGTGAAAAGTGTTACTTTATGAAATACAAATGAGGCTCTAAGCAATGCTTTTCGCTTACGCTTTTCGATAAGCGTACAAGTAGTTCCCCTACCGAAGGCCTATAAATGTGACTGCACATGTATCATCATAATTTGTTGATATACTTCGTTTATACCCGACTACGCATCGGCTAAAGCTGAGGGATCGGTGCACTATATAAGCTTCTCCTCGAAGTTCCCAAGCCACAAGTCGCTGATAATTCAAACAGAAATCATTTACCAAGCTCCGTGAGAACCTTTTCCAAT
